# Expression and role of Galectin-3 in the postnatal development of the cerebellum

**DOI:** 10.1101/364760

**Authors:** Inés González-Calvo, Fekrije Selimi

**Affiliations:** Center for Interdisciplinary Research in Biology (CIRB), Collège de France, CNRS UMR 7241, INSERM U1050, Paris, France

## Abstract

Many proteins initially identified in the immune system play roles in neurogenesis, neuronal migration, axon guidance, synaptic plasticity and other processes related to the formation and refinement of neural circuits. Although the function of the immune-related protein Galectin-3 (LGALS3) has been extensively studied in the regulation of inflammation, cancer and microglia activation, little is known about its role in the development of the brain. In this study, we identified that LGALS3 is expressed in the developing postnatal cerebellum. More precisely, LGALS3 is expressed by cells in meninges and in the choroid plexus, and in subpopulations of astrocytes and of microglial cells in the cerebellar cortex. Analysis of *Lgals3* knockout mice showed that *Lgals3* is dispensable for the development of cerebellar cytoarchitecture and Purkinje cell excitatory synaptogenesis in the mouse.

## Introduction

Many proteins initially identified in the immune system are also expressed in the developing and adult central nervous system (CNS) and contribute to a variety of processes during the formation and refinement of neural circuits. Immune system molecules play roles in neurogenesis, neuronal migration, axon guidance and synaptic plasticity (Reviewed in [1,2]). For example, the complement system proteins C1Q and C3, and the transmembrane major histocompatibility complex molecule class I (MHC-I) participate in the promotion of synapse elimination in the developing retinogeniculate pathway and in the vertebrate neuromuscular junction [3,4]. C1Q-related proteins such as the cerebellins and the C1Q-like proteins are known to be essential for synaptogenesis [5–8]. All these immune and immune-related proteins are expressed in neurons [3–8].

Galectins are a class of secreted lectins that play roles in various organs including the immune system. Galectins have been involved in inflammation, tumorigenesis, cancer, cell growth and metastasis [9–12]. They regulate cell migration, adhesion to the extracellular matrix, and cell-survival depending on their intra- or extracellular location [13]. The mammalian galectin family of soluble lectins is composed of fifteen members, all of which share a homologous carbohydrate recognition domain (CRD) that typically binds β-galactoside residues [14]. There are three architectural types of galectins: proto, tandem-repeat and chimera. The proto-galectins (Galectin-1, 2, 5, 7, −10, −11, −13, −14 and −15) present two homologous CRD (homodimers); the tandem-repeat-galectins (Galectin-4, 6, 8, −9 and −12) are composed by two distinct CRD (heterodimers). Both proto and tandem-repeat galectins have only carbohydrates as ligands. The pleiotropic LGALS3 is the only chimera type of galectins, and is particular in that in addition to the CRD domain, it contains a domain that enables its interaction with non-carbohydrate ligands [15] and the formation of pentamers and heterogeneous complexes with multivalent carbohydrates [16]. LGALS3 has been shown to interact with a broad collection of partners and to play roles at different time points during the life of the cell, including growth, adhesion, differentiation, cell-cycle and apoptosis [15]. Finally LGALS3 is involved in various pathologies such as cancer, inflammation and heart disease [17–19].

The role of LGALS3 in the central nervous system remain poorly understood. LGALS3 has been proposed to play a role in brain pathology. Its expression is increased in microglial cells upon various neuroinflammatory stimuli as, for instance, after ischemic injury [20,21]. LGALS3 binds the toll-like receptor 4 (TLR4) and promotes activation of microglia, a function that might promote the inflammatory response and neuronal death after ischemia [18]. The expression of LGALS3 also increases in reactive astrocytes in the injured cerebral cortex [19]. The role of LGALS3 in normal brain development and function has little been explored. LGALS3 is expressed in the subventricular zone, precisely in astrocytes around neuroblasts, and loss of LGALS3 reduces neuronal migration from the subventricular zone to the olfactory bulb *in vivo* [22]. LGALS3 plays also a role in oligodendrocyte differentiation and contributes to myelin function [23]. It has been shown that LGALS3 is not expressed in neurons in the olfactory bulb [22] or that it is only expressed in neurons under conditions of acute brain inflammation in the hippocampus [18] or in the injured cerebral cortex [19]. However, recently, LGALS3 has been reported to be expressed in hippocampal neurons and to play a role in memory formation through its interaction with integrin α3 [24]. LGALS3 can interact with the cell adhesion molecule integrin α3β1 [25], and integrin β1 induces the expression of LGALS3 *in vitro* [26]. While integrins have been involved in synaptogenesis and synapse modulation, the role of LGALS3 in these processes has not been explored so far. In this study, we investigated the expression and the role of *Lgals3* during development of the olivocerebellar network in the mouse. The connectivity and physiology of this neuronal network, and its development, have been well described, making it an ideal model to study the molecular mechanisms regulating the development of neuronal networks. Our data show that while LGALS3 is expressed during postnatal development in the olivocerebellar network, it is mainly expressed by glial cells, and it is dispensable for neuronal development and synaptogenesis in this structure.

## Material and methods

### Ethics Statement

All animal protocols were approved by the *Comité Regional d’Ethique en Experimentation Animale* (# 00057.01).

### Animals

*Lgals3*^+/+^ wild-type and *Lgals3*^-/-^ knockout littermates were obtained by breeding heterozygotes (129Sv background) and were kindly provided by Dr. Françoise Poirier [27]. CX_3_CR1^eGFP/eGFP^ mice were kindly provided by Dr. Etienne Audinat and Prof. S. Jung [28].

### Antibodies

The following primary antibodies were used: mouse monoclonal anti-CABP (1:1000; swant, Cat#300), rabbit polyclonal anti-CABP (1:1000; swant, Cat#9.03), mouse monoclonal anti-GFAP (1:500; millipore, Cat#MAB360), rabbit polyclonal anti-GLURδ1/2 (1:1000; millipore, Cat#AB2285), goat polyclonal anti-LGALS3 (1:200; R&D Systems, Cat#AF1197), mouse monoclonal anti-OLIG2 (1:500; millipore, Cat#MABN50), guinea pig polyclonal anti-VGLUT1 (1:5000; millipore, Cat#AB5905) and guinea pig polyclonal anti-VGLUT2 (1:5000; millipore, Cat#AB2251). The following secondary antibodies were used: donkey polyclonal anti-goat Alexa Fluor 568 (1:1000; invitrogen, Cat#A11057), donkey polyclonal anti-mouse Alexa Fluor 488 (1:1000; invitrogen, Cat#R37114), donkey polyclonal anti-rabbit Alexa Fluor 488 (1:1000; invitrogen, Cat#A21206) and goat polyclonal anti-guinea pig Alexa Fluor 594 (1:1000; Invitrogen, Cat#A11076).

### RTqPCR

RNA samples were obtained from cerebellar and brainstem tissues using the RNeasy Mini kit (QIAGEN, Hilden, Germany) and cDNA amplified using the SuperScript^®^ VILO ^™^ cDNA Synthesis kit (life technologies, Paisley, UK) according to manufacturer’s instructions. Quantitative PCR was performed using the TaqMan Universal Master Mix II with UNG (Applied Biosystems, Courtboeuf, France) and the following TaqMan probes: *Rpl13a* (#4331182_Mm01612986_gH) and *Lgals3* (#4331182_ Mm00802901_m1).

### Immunohistochemistry

Immunostaining was performed on 30μm-thick parasagittal sections obtained using a freezing microtome from brains of mice perfused with 4% paraformaldehyde in phosphate buffered saline (PBS) solution. Sections were washed three times for five minutes in PBS, then blocked with PBS 4% donkey serum (DS; abcam, Cat#ab7475) for 30 minutes. The primary antibodies were diluted in PBS, 1% DS, 1% Triton X-100 (Tx; sigma-aldrich, Cat#x100). The sections were incubated in the primary antibody solution overnight at 4°C and then washed three times for five minutes in PBS 1%Tx. Sections were incubated in the secondary antibody, diluted in PBS 1%DS 1%Tx solution, for 1h at room temperature. The sections were then incubated for 15 minutes with the nuclear marker Hoechst 33342 (sigma-aldrich, Cat#H6024), followed by three washes for five minutes in PBS 1%Tx and recovered in PBS. The sections were mounted with Prolong Gold (invitrogen; Cat#P36960).

### Image acquisition and analysis

Imaging was performed using confocal microscopy (SP5, leica). The pinhole aperture was set to 1 Airy Unit and a z-step of 500 nm was used. The software ImageJ was used to measure the area of the cerebellum from images of staining obtained with the nuclear marker Hoechst, and the area and the length of the molecular layer using images of the anti-CABP staining.

### Statistical Analysis

Data generated with ImageJ were imported in GraphPad Prism for statistical analysis. Values are given as mean ± s.e.m. Normality was assessed using the Kolmogorov-Smirnov normality test. Differences between two groups were tested using the two-tailed Student’s t-test. Groups were considered significantly different when at least a 95% confidence level (P<0.05) was obtained.

## Results

### Lgals3 mRNA is expressed in the cerebellum and brainstem during postnatal development

The expression level of *Lgals3* mRNA in the cerebellum was measured using quantitative RT-PCR at different time points during late embryonic and postnatal development. The expression of *Lgals3* mRNA is developmentally regulated in the cerebellum (Fig 1A). It peaks at P14 when it quadruples compared to the levels detected at E17 and P0 and returns to the levels detected during the first postnatal week by P21, a stage when neurogenesis, differentiation and synaptogenesis in the cerebellum is to a large extent complete [29]. This developmentally regulated expression profile is similar to the mRNA expression profile of the adhesion G protein coupled receptor BAI3, that was recently shown to regulate Purkinje cell synaptogenesis [5,7]. This pattern thus suggested a role for LGALS3 during cerebellar Purkinje cell development and excitatory synaptogenesis.

**Fig 1.**
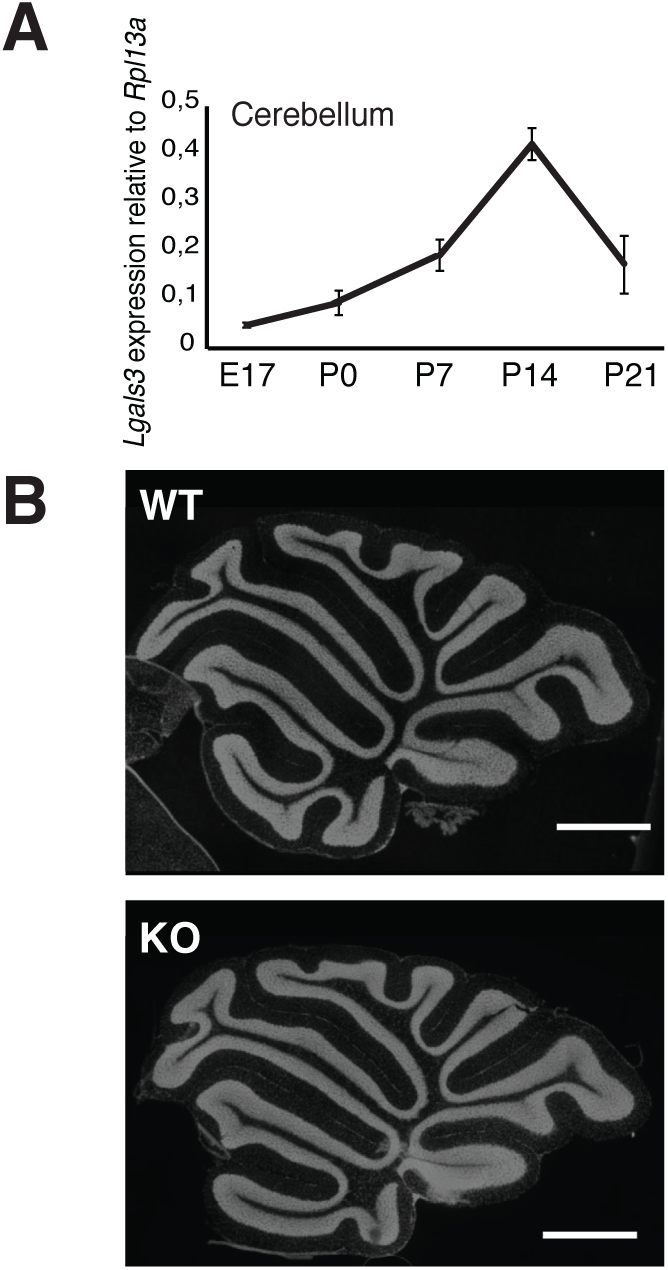
The cerebellar architecture is not affected in *Lgals3* null mice. (A) *Lgals3* mRNA expression relative to Rpl13a measured by RTqPCR in the cerebellum during development. E17: embryonic day 17. P0, P7, P14, and P21: postnatal day 0, 7, 14, and 21. (B) *Lgals3* knockout (KO) mice present no defects in cerebellar morphology. Parasagittal cerebellar slices of wild-type (WT) and *Lgals3* KO adult mice were stained using the nuclear marker Hoechst.

### Cerebellar architecture and excitatory synaptogenesis are not affected by Lgals3 invalidation

To study the role of LGALS3 in the development of the cerebellum, we analyzed mice invalidated for the *Lgals3* gene [27]. The general architecture of the cerebellum, the different layers and folia, was normal in those mice (Fig 1B). Quantitative analysis showed a slight decrease (less than 10%) in the mean area of the cerebellum and in the mean area and length of the molecular layer in adult *Lgals3* knockout (KO) mice when compared to *Lgals3* wildtype (WT) mice (mean cerebellar area: 10.39×10^6^±1.72×10^5^ μm^2^ versus 9.62×10^6^±2.34×10^5^ μm^2^, P=0.0235; mean molecular layer area: 4.9×10^6^±0.81×10^5^ μm^2^ versus 4.42×10^6^±0.72×10^5^ μm^2^, P=0.0015; mean molecular layer length: 115.95±1.65 μm versus 105.5±1.53 μm, P=0.000082; for WT and KO six months-old mice, respectively; Student’s t-test; n=3 animals per genotype). These small differences were associated with a decrease in body weight in adult KO (mean body weight ± s.e.m.: 26.033±0.07 g for WT, 24.003±0.66 g for KO; n=5 animals per genotype, P=0.027, Student’s t-test). These results show that lack of LGALS3 does not have a major effect on neurogenesis and neuronal morphogenesis in the cerebellum. *Lgals3* expression peaks at postnatal day 14 (Fig 1A), a period of intense synaptogenesis in the cerebellar cortex, and LGALS3 potentially interacts with integrin β1 that is present at the Climbing fiber (CF)-Purkinje cell synapse [25,30]. To analyze the possible role of LGALS3 in excitatory synapse formation we used immunolabeling with the pre-synaptic marker VGLUT2, to label the CF presynaptic boutons, and with the pre-synaptic marker VGLUT1, specific for the parallel fiber (PF) presynaptic boutons in the cerebellar cortex. At P15, VGLUT2 clusters are found on the somato-dendritic region of Purkinje cells, in particular on proximal dendrites both in *Lgals3* KO and *Lgals3* WT mice (Fig 2A). Smaller VGLUT2 clusters corresponding to maturing PF-Purkinje cell synapses [31] are found in distal dendrites, in particular in the upper part of the molecular layer, in both genotypes. In adult mice, when CF and PF synapses are mature, VGLUT2 clusters are only found on the Purkinje cells proximal dendrites and extend to about 4/5^th^ of the molecular layer height both in *Lgals3* KO and WT cerebella (Fig 2B). The pattern of VGLUT1 immunostaining is similar in *Lgals3* KO mice and control littermates, with a sparse staining of distal dendrites at P15 (Fig 2C) and a dense staining non-overlapping with the VGLUT2 clusters at P65 (Fig 2D). These qualitative data show that both PF and CF excitatory synaptogenesis in cerebellar Purkinje cells is not affected by *Lgals3* invalidation.

**Fig 2.**
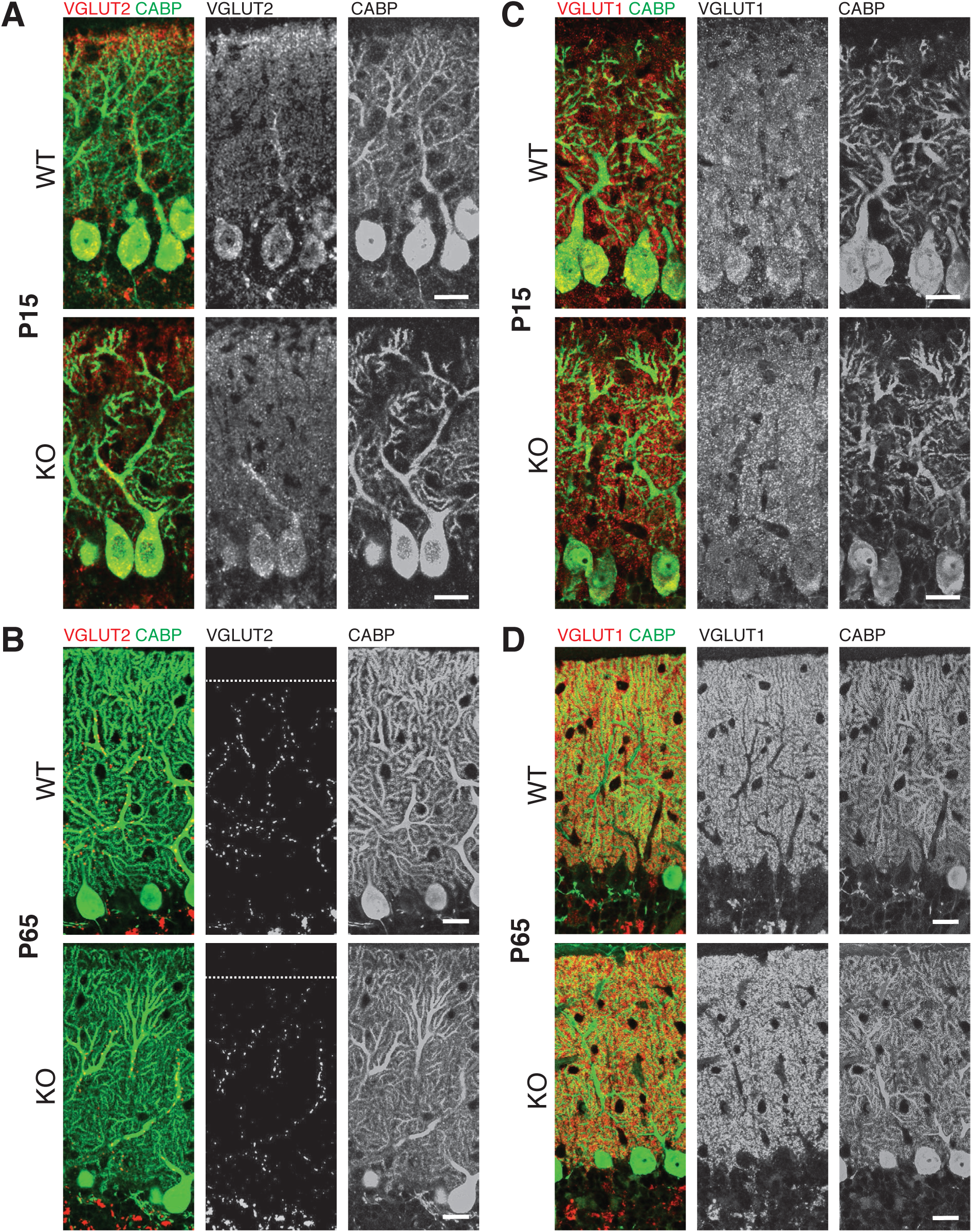
No changes in excitatory Purkinje cell synapses in *Lgals3* knockout mice. (A and B) Immunostaining for the Purkinje cell marker CABP (green) and the presynaptic marker VGLUT2 (red) was performed on sections from *Lgals3* WT and KO mice at P15 (A) and at P65 (B). (C and D) Immunostaining for the Purkinje cell marker CABP (green) and the presynaptic marker VGLUT1 (red) specific for mature parallel fiber synapses was performed on sections from *Lgals3* WT and KO mice at P15 (C) and at P65 (D). Scale bar: 25μm. Data are representative of three independent experiments.

### LGALS3 is expressed in glia in the cerebellum

The lack of effect on cerebellar morphology and synaptogenesis of *Lgals3* invalidation raised the question as to what type of cells express the LGALS3 protein during postnatal development. Using a LGALS3 antibody, we localized LGALS3 expression in the cerebellum and brainstem during postnatal development. The specificity of the antibody was confirmed by the lack of immunoreactivity in cerebellar sections from *Lgals3* KO mice (n=3 independent experiments; Fig 3A). At P15, the highest expression of LGALS3 was found in the choroid plexus located above the cerebellum, and in the ependymal cells and meningeal cells lining the cerebral aqueduct and the 4^th^ ventricle. In the cerebellar cortex, the white matter is strongly stained, while rare scattered cells in the grey matter also display some LGALS3 expression. Despite a more than two-fold decrease in *Lgals3* mRNA levels in the cerebellum between P14 and P21 (Fig 1A), the pattern of LGALS3 immunostaining was similar at P15 and P22 (Fig 3B and C respectively). At P15, Purkinje cells display a staining that is barely distinguishable from background, and that completely disappears at P21. In the brainstem, LGALS3 immunostaining was mainly found in the white matter underneath the 4^th^ ventricle, between the pons (medial vestibular nucleus) and the medulla (pontine central grey) at both P15 and P22 (Figs 3B-C).

**Fig 3.**
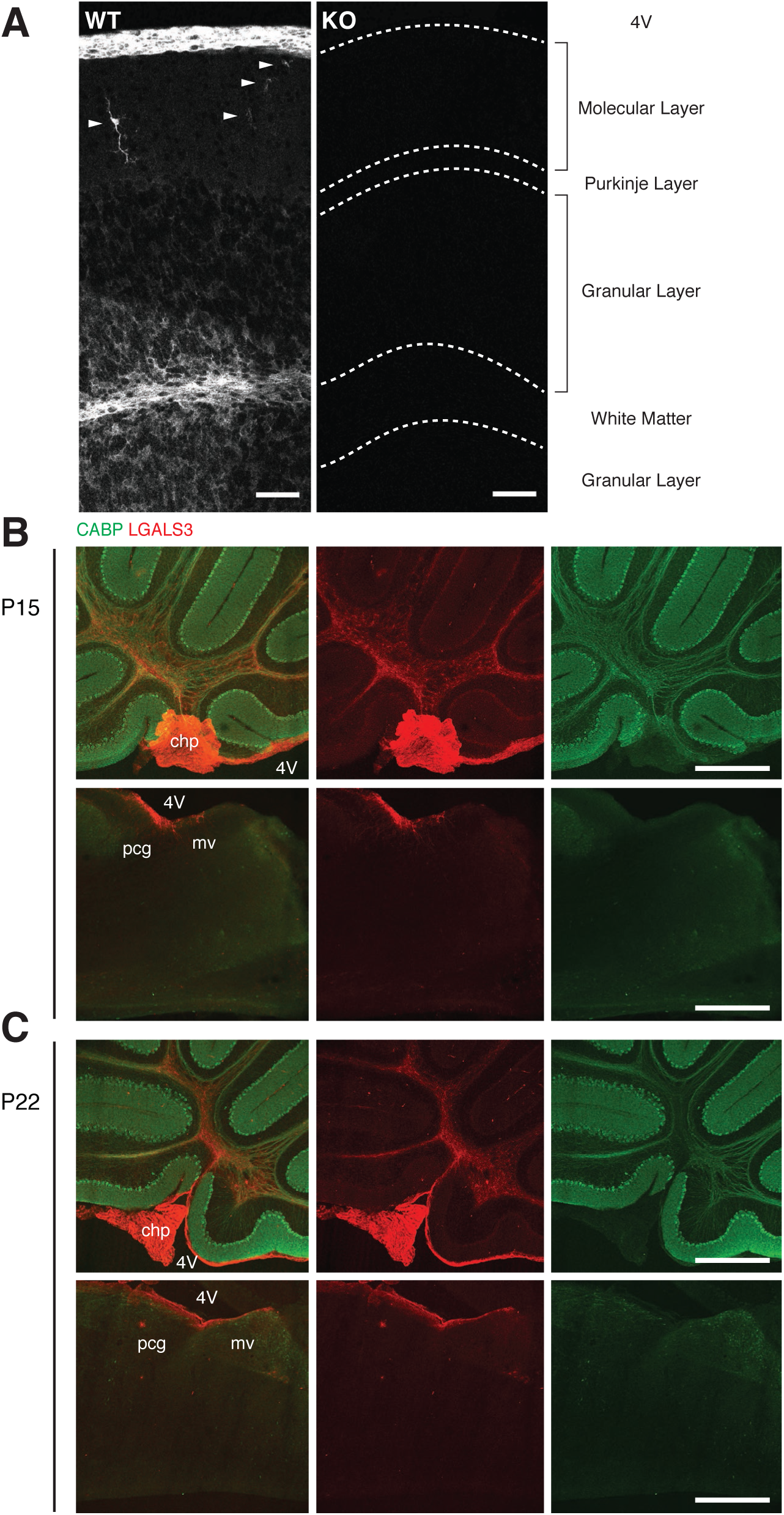
LGALS3 expression in the cerebellum and brainstem during postnatal development. (A) Immunohistochemistry for LGALS3 show the specificity of the LGALS3 antibody. WT: wild-type mice; KO: knockout mice. Positive cells in the molecular layer are marked with a white arrowhead, scale bar 50μm. (B) Immunohistochemistry for the Purkinje cell marker CABP (green) and LGALS3 (red) on parasagittal cerebellar sections from P15 and P22 wild-type mice. Scale bar=100μm. Data are representative of three independent experiments. Legends: 4V, 4^th^ ventricle; chp, choroid plexus; pcg, pontine central gray and mv, medial vestibular nucleus.

The cerebellar white matter contains both oligodendrocytes and astrocytes. Previous studies have detected LGALS3 expression in Schwann cells of the peripheral nervous system [32] and in oligodendrocytes at different stages of maturation [23]. Co-immunostaining experiments using an antibody against the oligodendrocyte specific marker OLIG-2 show some LGALS3 expression in OLIG-2 positive oligodendrocytes (Fig 4A), while co-immunostaining experiments with an anti-GFAP antibody to label astrocytes showed extensive colocalization of LGALS3 and GFAP in cells of the cerebellar white matter (Fig 4B). In the grey matter of the cerebellum, the morphology of scattered LGALS3 positive cells was reminiscent of microglial cells. Immunolabeling of cerebellar sections from a CX_3_CR1^eGFP/eGFP^ mouse line that expresses soluble GFP in microglial cells [28], revealed double labeled cells and confirmed the microglial identity of LGALS3 positive cells in the molecular layer (Fig 4C). Overall our study shows that LGALS3 is expressed in non-neuronal cells in the cerebellum and brainstem during postnatal development, in particular subpopulations of astrocytes and microglial cells.

**Fig 4.**
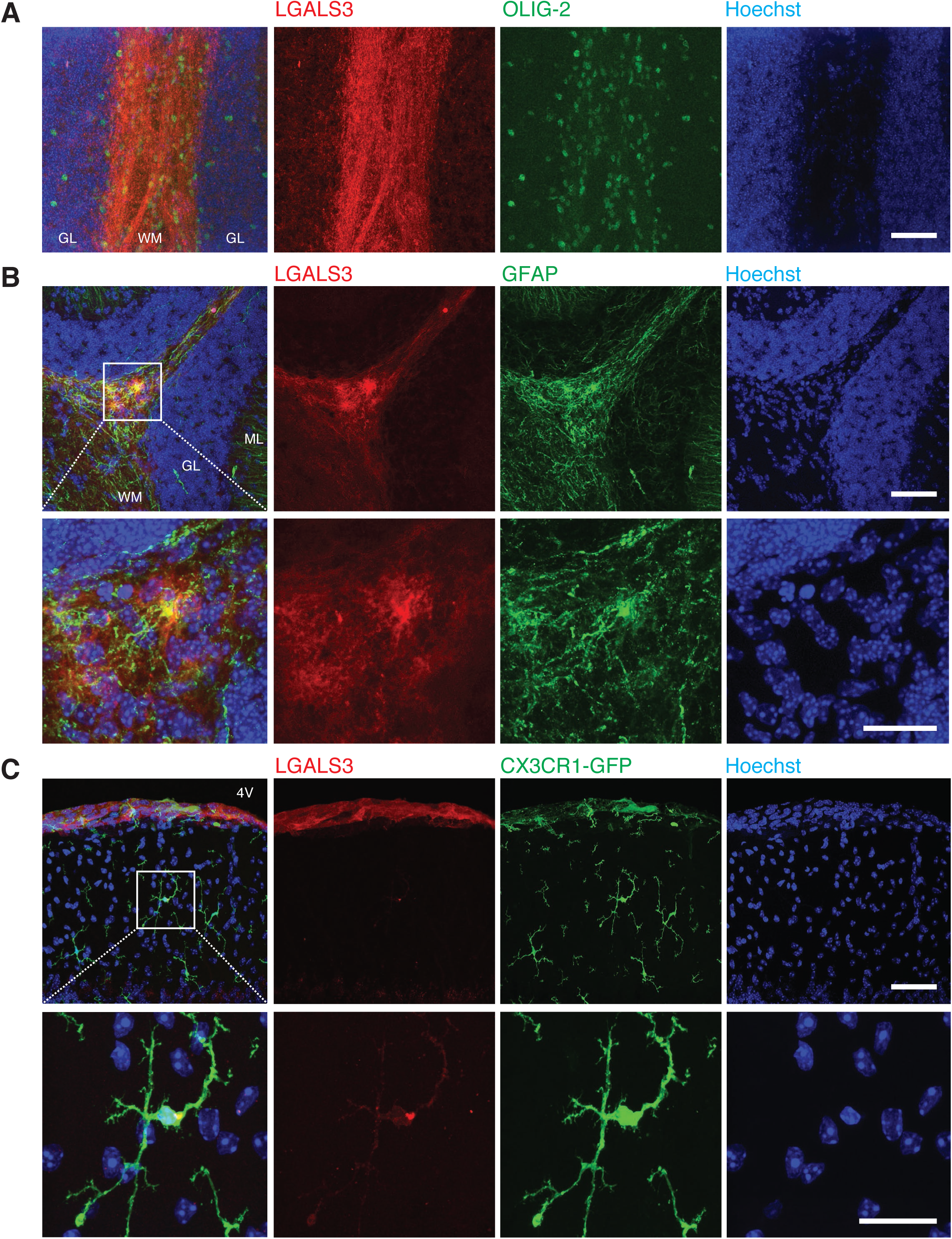
LGALS3 protein expression in meningeal cells and in glial cells of the cerebellar cortex of the adult mice. (A) LGALS3 expression in oligodendrocytes in the cerebellar white matter. Co-immunostaining of parasagittal cerebellar slices using anti-LGALS3 (red), oligodendrocyte marker anti-OLIG-2 (green) and the nuclear marker Hoechst (blue). (B) LGALS3 expression in astrocytes in the cerebellar white matter. Co-immunostaining of parasagittal cerebellar slices using anti-LGALS3 (red), astrocyte marker anti-GFAP (green) and the nuclear marker Hoechst (blue). (C) LGALS3 expression in some microglial cells of the molecular layer. Immunostaining anti-LGALS3 (red) in parasagittal sections of mice CX_3_CR1^eGFP/eGFP^, microglial CX_3_CR1-GFP (green) and the nuclear marker Hoechst (blue). Scale bar 50μm and 20μm for the magnification. Data are representative of three independent experiments. Legends: 4V, 4^th^ ventricle; GL, granular layer; ML, molecular layer and WM, white matter.

## Discussion

The role of LGALS3 in the development of the CNS remains poorly studied. We show that during postnatal development, LGALS3 is expressed in the choroid plexus in the fourth ventricle, as well as in a subset of glial cells in the cerebellar cortex. This expression is dispensable for the development of the normal cytoarchitecture of the cerebellum and excitatory synaptogenesis in Purkinje cells.

LGALS3 expression has been described previously in pathological conditions in the mammalian brain, in particular after ischemia [20,21]. LGALS3 is expressed by a subset of proliferating astrocytes that contribute to reactive gliosis [19]. In the normal brain, high expression of LGALS3 has been found in the subventricular zone, rostral migratory stream and olfactory bulb [22] and in the subependymal zone of the lateral ventricles [19]. Our results are in concordance, showing high expression of LGALS3 in the choroid plexus beneath the cerebellum and in the subependymal zone of the fourth ventricle. In the brain parenchyma, some studies have reported neuronal expression of LGALS3 in the hippocampus and cerebellum [24,33,34], while others did not detect any neuronal expression [19,35]. In the cerebellum in particular, low levels of mRNA expression were reported in Purkinje cells and granule cells in an analysis of Allen Brain Atlas data by John and Mishra [33]. Our results showed very low to undetectable expression of LGALS3 in cerebellar neurons, in agreement with previous results [35,36]. However high levels of LGALS3 were found in subsets of astrocytes and subsets of microglia in the cerebellar cortex. This is in accordance with previous results showing expression of LGALS3 by microglia [18] and astrocytes *in vitro* [23]. LGALS3 has been involved in myelination [23]. Our results show some expression of LGALS3 in oligodendrocytes of the cerebellar white matter during postnatal development. Together with its expression in astrocytes of the white matter, this suggests an implication of LGALS3 in the crosstalk between astrocytes and oligodendrocytes during myelination (reviewed in [37]).

Several reasons suggested a role for *Lgals3* during synaptogenesis. First, LGALS3 is a secreted and immune-related protein with a developmentally regulated mRNA expression profile that coincides with the timing of excitatory synaptogenesis in the cerebellum. These characteristics are shared by C1QL1 and CBLN1, two major synaptogenic proteins of the CF-Purkinje cell and PF-Purkinje cell synapses respectively [5–8][5–8]. Second, LGALS3 promotes neural cell adhesion and neurite outgrowth in cultured neurons [38]. Third it has been shown that LGALS3 interacts functionally with integrins, adhesion proteins known to play roles in synaptogenesis and synapse modulation. Integrin α3β1 is necessary for the organization of the cerebellar excitatory synapse CF-Purkinje cell [30], LGALS3 can interact with integrin α3β1 [25], and integrin β1 induces the expression of LGALS3 in culture [26]. Our analysis of the *Lgals3* knockout mouse did not find any major perturbation of Purkinje cell excitatory synaptogenesis in the absence of LGALS3. Our data thus indicate that either LGALS3 plays no role in synaptogenesis, or that other proteins play redundant functions. Our results do not exclude a role for LGALS3 in regulating synapse maturation and function. In particular, microglia have been involved in synapse elimination during development of the visual system [3], and LGALS3 promotes phagocytosis of PC12 cells by microglia [39]. Activity-dependent synapse refinement of the CF-Purkinje cell synapse happens largely during the first three postnatal weeks [29]. Given LGALS3 expression in microglial cells in the cerebellar cortex during this period, it would be interesting to test whether LGALS3 is involved in climbing fiber synapse elimination, in particular, through its interaction with integrin α3β1, a known regulator of CF synaptogenesis.

## Author contributions

F.S. and I.G.C. designed the experiments and wrote the manuscript. I.G.C. performed the experiments.

## Acknowledgments

We thank Dr. F. Poirier for *Lgals3*^+/+^ and *Lgals3*^-/-^ mice, Prof. S. Jung and Dr. E. Audinat for CX_3_CR1^eGFP/eGFP^ mice, and Maëva Talleur for the RTqPCR. We also thank the CIRB Imaging Facility.

This work has received support under the Investissement d’Avenir program launched by the French government and implemented with the ANR-11-IDEX-0001-02 PSL* Research University (to I.G.C. and F.S.). Funding was also provided by the ATIP AVENIR program (to F.S.) and Fondation pour la Recherche Medicale (DEQ20150331748).

## References

1. Boulanger LM. Immune proteins in brain development and synaptic plasticity. Neuron. 2009;64: 93–109. doi:10.1016/j.neuron.2009.09.001

2. Carpentier PA, Palmer TD. Immune Influence on Adult Neural Stem Cell Regulation and Function. Neuron. 2009;64: 79–92. doi:10.1016/j.neuron.2009.08.038

3. Stevens B, Allen NJ, Vazquez LE, Howell GR, Christopherson KS, Nouri N, et al. The classical complement cascade mediates CNS synapse elimination. Cell. 2007;131: 1164–1178. doi:10.1016/j.cell.2007.10.036

4. Tetruashvily MM, McDonald MA, Frietze KK, Boulanger LM. MHCI promotes developmental synapse elimination and aging-related synapse loss at the vertebrate neuromuscular junction. Brain Behav Immun. 2016;56: 197–208. doi:10.1016/j.bbi.2016.01.008

5. Kakegawa W, Mitakidis N, Miura E, Abe M, Matsuda K, Takeo YH, et al. Anterograde C1ql1 signaling is required in order to determine and maintain a single-winner climbing fiber in the mouse cerebellum. Neuron. 2015;85: 316–329. doi:10.1016/j.neuron.2014.12.020

6. Matsuda K, Miura E, Miyazaki T, Kakegawa W, Emi K, Narumi S, et al. Cbln1 is a ligand for an orphan glutamate receptor delta2, a bidirectional synapse organizer. Science. 2010;328: 363–368. doi:10.1126/science.1185152

7. Sigoillot SM, Iyer K, Binda F, González-Calvo I, Talleur M, Vodjdani G, et al. The Secreted Protein C1QL1 and Its Receptor BAI3 Control the Synaptic Connectivity of Excitatory Inputs Converging on Cerebellar Purkinje Cells. Cell Rep. 2015;10: 820–832. doi:10.1016/j.celrep.2015.01.034

8. Uemura T, Lee S-J, Yasumura M, Takeuchi T, Yoshida T, Ra M, et al. Trans-synaptic interaction of GluRdelta2 and Neurexin through Cbln1 mediates synapse formation in the cerebellum. Cell. 2010;141: 1068–1079. doi:10.1016/j.cell.2010.04.035

9. Dumic J, Dabelic S, Flögel M. Galectin-3: an open-ended story. Biochim Biophys Acta. 2006;1760: 616–635. doi:10.1016/j.bbagen.2005.12.020

10. Liu F-T, Rabinovich GA. Galectins: regulators of acute and chronic inflammation. Ann N Y Acad Sci. 2010;1183: 158–182. doi:10.1111/j.1749-6632.2009.05131.x

11. Nangia-Makker P, Nakahara S, Hogan V, Raz A. Galectin-3 in apoptosis, a novel therapeutic target. J Bioenerg Biomembr. 2007;39: 79–84. doi:10.1007/s10863-006-9063-9

12. Ochieng J, Furtak V, Lukyanov P. Extracellular functions of galectin-3. Glycoconj J. 2004;19: 527–535. doi:10.1023/B:GLYC.0000014082.99675.2f

13. Elola MT, Wolfenstein-Todel C, Troncoso MF, Vasta GR, Rabinovich GA. Galectins: matricellular glycan-binding proteins linking cell adhesion, migration, and survival. Cell Mol Life Sci CMLS. 2007;64: 1679–1700. doi:10.1007/s00018-007-7044-8

14. Leffler H, Carlsson S, Hedlund M, Qian Y, Poirier F. Introduction to galectins. Glycoconj J. 2004;19: 433–440. doi:10.1023/B:GLYC.0000014072.34840.04

15. Barondes SH, Cooper DN, Gitt MA, Leffler H. Galectins. Structure and function of a large family of animal lectins. J Biol Chem. 1994;269: 20807–20810.

16. Ahmad N, Gabius H-J, André S, Kaltner H, Sabesan S, Roy R, et al. Galectin-3 precipitates as a pentamer with synthetic multivalent carbohydrates and forms heterogeneous cross-linked complexes. J Biol Chem. 2004;279: 10841–10847. doi:10.1074/jbc.M312834200

17. Wesley UV, Vemuganti R, Ayvaci R, Dempsey RJ. Galectin-3 enhances angiogenic and migratory potential of microglial cells via modulation of integrin linked kinase signaling. Brain Res. 2013;1496: 1–9. doi:10.1016/j.brainres.2012.12.008

18. Burguillos MA, Svensson M, Schulte T, Boza-Serrano A, Garcia-Quintanilla A, Kavanagh E, et al. Microglia-Secreted Galectin-3 Acts as a Toll-like Receptor 4 Ligand and Contributes to Microglial Activation. Cell Rep. 2015;10: 1626–1638. doi:10.1016/j.celrep.2015.02.012

19. Sirko S, Irmler M, Gascón S, Bek S, Schneider S, Dimou L, et al. Astrocyte reactivity after brain injury-: The role of galectins 1 and 3. Glia. 2015;63: 2340–2361. doi:10.1002/glia.22898

20. Satoh K, Niwa M, Binh NH, Nakashima M, Kobayashi K, Takamatsu M, et al. Increase of galectin-3 expression in microglia by hyperthermia in delayed neuronal death of hippocampal CA1 following transient forebrain ischemia. Neurosci Lett. 2011;504: 199–203. doi:10.1016/j.neulet.2011.09.015

21. Satoh K, Niwa M, Goda W, Binh NH, Nakashima M, Takamatsu M, et al. Galectin-3 expression in delayed neuronal death of hippocampal CA1 following transient forebrain ischemia, and its inhibition by hypothermia. Brain Res. 2011;1382: 266–274. doi:10.1016/j.brainres.2011.01.049

22. Comte I, Kim Y, Young CC, van der Harg JM, Hockberger P, Bolam PJ, et al. Galectin-3 maintains cell motility from the subventricular zone to the olfactory bulb. J Cell Sci. 2011;124: 2438–2447. doi:10.1242/jcs.079954

23. Pasquini LA, Millet V, Hoyos HC, Giannoni JP, Croci DO, Marder M, et al. Galectin-3 drives oligodendrocyte differentiation to control myelin integrity and function. Cell Death Differ. 2011;18: 1746–1756. doi:10.1038/cdd.2011.40

24. Chen Y-C, Ma Y-L, Lin C-H, Cheng S-J, Hsu W-L, Lee EH-Y. Galectin-3 Negatively Regulates Hippocampus-Dependent Memory Formation through Inhibition of Integrin Signaling and Galectin-3 Phosphorylation. Front Mol Neurosci. 2017;10: 217. doi:10.3389/fnmol.2017.00217

25. Saravanan C, Liu F-T, Gipson IK, Panjwani N. Galectin-3 promotes lamellipodia formation in epithelial cells by interacting with complex N-glycans on alpha3beta1 integrin. J Cell Sci. 2009;122: 3684–3693. doi:10.1242/jcs.045674

26. Margadant C, van den Bout I, van Boxtel AL, Thijssen VL, Sonnenberg A. Epigenetic Regulation of Galectin-3 Expression by β1 Integrins Promotes Cell Adhesion and Migration. J Biol Chem. 2012;287: 44684–44693. doi:10.1074/jbc.M112.426445

27. Colnot C, Fowlis D, Ripoche MA, Bouchaert I, Poirier F. Embryonic implantation in galectin 1/galectin 3 double mutant mice. Dev Dyn Off Publ Am Assoc Anat. 1998;211: 306–313. doi:10.1002/(SICI)1097-0177(199804)211:4<306::AID-AJA2>3.0.C0;2-L

28. Jung S, Aliberti J, Graemmel P, Sunshine MJ, Kreutzberg GW, Sher A, et al. Analysis of fractalkine receptor CX(3)CR1 function by targeted deletion and green fluorescent protein reporter gene insertion. Mol Cell Biol. 2000;20: 4106–4114.

29. Hashimoto K, Kano M. Synapse elimination in the developing cerebellum. Cell Mol Life Sci CMLS. 2013;70: 4667–4680. doi:10.1007/s00018-013-1405-2

30. Su J, Stenbjorn RS, Gorse K, Su K, Hauser KF, Ricard-Blum S, et al. Target-derived matricryptins organize cerebellar synapse formation through α3β1 integrins. Cell Rep. 2012;2: 223–230. doi:10.1016/j.celrep.2012.07.001

31. Hashimoto K, Ichikawa R, Kitamura K, Watanabe M, Kano M. Translocation of a “winner” climbing fiber to the Purkinje cell dendrite and subsequent elimination of “losers” from the soma in developing cerebellum. Neuron. 2009;63: 106–118. doi:10.1016/j.neuron.2009.06.008

32. Reichert F, Saada A, Rotshenker S. Peripheral nerve injury induces Schwann cells to express two macrophage phenotypes: phagocytosis and the galactose-specific lectin MAC-2. J Neurosci Off J Soc Neurosci. 1994;14: 3231–3245.

33. John S, Mishra R. mRNA Transcriptomics of Galectins Unveils Heterogeneous Organization in Mouse and Human Brain. Front Mol Neurosci. 2016;9: 139. doi:10.3389/fnmol.2016.00139

34. Yoo H-I, Kim E-G, Lee E-J, Hong S-Y, Yoon C-S, Hong M-J, et al. Neuroanatomical distribution of galectin-3 in the adult rat brain. J Mol Histol. 2017;48: 133–146. doi:10.1007/s10735-017-9712-9

35. Kobayashi K, Niwa M, Hoshi M, Saito K, Hisamatsu K, Hatano Y, et al. Early microlesion of viral encephalitis confirmed by galectin-3 expression after a virus inoculation. Neurosci Lett. 2015;592: 107–112. doi:10.1016/j.neulet.2015.02.061

36. Mahoney SA, Wilkinson M, Smith S, Haynes LW. Stabilization of neurites in cerebellar granule cells by transglutaminase activity: identification of midkine and galectin-3 as substrates. Neuroscience. 2000;101: 141–155.

37. Domingues HS, Portugal CC, Socodato R, Relvas JB. Oligodendrocyte, Astrocyte, and Microglia Crosstalk in Myelin Development, Damage, and Repair. Front Cell Dev Biol. 2016;4: 71. doi:10.3389/fcell.2016.00071

38. Pesheva P, Kuklinski S, Schmitz B, Probstmeier R. Galectin-3 promotes neural cell adhesion and neurite growth. J Neurosci Res. 1998;54: 639–654.

39. Nomura K, Vilalta A, Allendorf DH, Hornik TC, Brown GC. Activated Microglia Desialylate and Phagocytose Cells via Neuraminidase, Galectin-3, and Mer Tyrosine Kinase. J Immunol Baltim Md 1950. 2017;198: 4792–4801. doi:10.4049/jimmunol.1502532

